# Enhancement of mRNA translation efficiency through 5’-UTR engineering

**DOI:** 10.1101/2025.09.30.679448

**Authors:** Esther Broset, Irene Blasco-Machín, Verónica Lampaya, Carlos Matute, Alfonso Toro-Córdova, Juan Martínez-Oliván

## Abstract

The rapid progress of mRNA therapeutics has underscored a persistent challenge: achieving high protein expression at low dose. The 5’ untranslated region (5’-UTR) is a key regulator of translation initiation efficiency, prompting the question of whether a simple and portable modification could enhance expression across diverse designs. Here, we systematically engineered short repeats of the Kozak “core” motif (5’-GCCACC-3’) immediately upstream of the start codon and evaluated constructs incorporated into two widely used human UTRs (APO and HBB) in HeLa and HEK293T cells. Translation enhancement displayed a non-monotonic dependence on the number of Kozak repeats, with three repetitions consistently outperforming the native sequence and any other configuration.

In mice, intramuscular lipid nanoparticle delivery of the three-copy design increased luciferase expression by ∼4-fold relative to the wild-type context and by up to ∼23-fold compared to a licensed vaccine UTR benchmark, providing clear in vivo relevance.

These findings demonstrate that fine-tuning AUG-proximal Kozak elements constitutes a broadly applicable, UTR-independent strategy to enhance translation efficiency, offering a simple yet powerful principle for dose-sparing mRNA design in therapeutic applications.

## Introduction

In recent years, mRNA vaccines and gene replacement therapies, which enable the exogenous production of proteins either to elicit an immune response or to compensate for defective genes, respectively, have significantly influenced clinical practice. In 2023, approximately 300 mRNA-based drugs were undergoing clinical trials (1), and by 2025 this number had risen to 854 (2), representing an exponential increase from fewer than 20 in 2019 (3), indicative of the rise in interest and the potential surrounding this class of drugs. Although mRNA represents a highly promising therapeutic strategy, there are still some aspects that can be improved. Specifically, augmenting the amount of protein expressed per mRNA molecule could lower the required dosage, thereby reducing manufacturing costs and potentially enhancing clinical outcomes.

Multiple strategies are being employed to enhance the potency of mRNA therapeutics, including improving delivery and cell transduction (4), increasing mRNA stability (5), optimizing regulatory sequences to boost translation efficiency. While strategies to improve delivery using lipid nanoparticles (LNPs) have received considerable attention (6), comparatively few efforts have focused on enhancing translation through the modulation of mRNA regulatory elements. In this study, we specifically address this gap by focusing on the optimization of 5’untranslated regions (5’-UTRs).

The 5’-UTR is recognized as one of the key determinants of mRNA translation efficiency; however, its complete underlying mechanisms remain incompletely understood. Several studies have screened 5’-UTR sequences and have related them to different levels of protein expression (7–11). Additionally, several elements have been described to affect 5’-UTR efficiency including upstream start codons and open reading frames (ORFs) (12), the presence of stable secondary structures (13) or 5’-Terminal Oligo Pyrimidine (5’TOP) motifs located within four nucleotides of the transcription start site (14). More recently, the influence of the modified N-1 methyl pseudouridine (m^1^Ψ), commonly used in mRNA therapeutics, on 5’-UTR translation efficiency has been investigated (15), enabling the identification of sequences that achieve optimal performance in the presence of this modification. Moreover, language models and deep learning approaches have been applied to predict and design highly efficient 5’-UTRs (16, 17). However, despite these advances, a clear framework for the rational design of improved 5’UTRs has yet to be established, and most current mRNA therapeutics continue to rely on 5’UTRs derived from highly expressed endogenous genes (18–21).

The structure and the cellular mechanism underlying translation initiation should also be considered when designing an efficient 5’-UTR (13). Once the mRNA reaches the cytosol, the CAP structure is recognized by eIF4E, which recruits eIF4G, while the poly(A) tail is bound by PABP, promoting mRNA circularization and assembly of the initiation complex. The 40S ribosomal subunit is then recruited and scans along the 5’-UTR until the start codon is identified. Subsequently, the 60S subunit joins the 40S to form the functional 80S ribosome, initiating the elongation phase. Based on this mechanism, an aptamer that binds the translation factor eIF4G has been proposed as a universal strategy for improving mRNA-based therapies (22). However, the presence strong secondary structures within the 5’-UTR should be carefully considered, as these can obstruct ribosomal scanning and block translation (23) or cause ribosomal go backtracking, thereby delaying initiation (24).

A pivotal regulatory element located within the 5’-UTR is the Kozak sequence (25, 26), a highly conserved consensus sequence in vertebrates positioned upstream of the AUG start codon (GCCRCC-AUGG). This sequence is critical for efficient ribosomal scanning and initiation of translation. The Kozak consensus sequence plays an essential role in proteomic regulation by influencing translation initiation efficiency and fidelity (27). Additionally, the Kozak motif, along with its flanking sequences, has been extensively engineered to fine-tune translational regulation, including the modulation of bidirectional translation (28).

Because the Kozak consensus sequence is the principal cis-regulatory element guiding the transition of the scanning pre-initiation complex into a committed initiation complex at the cognate AUG, we hypothesized that a short “Kozak ramp”, consisting of a tandem repeat of the core GCCACC motif placed immediately upstream of the start codon, could improve start site recognition and enhance protein yield. Extending Kozak-like sequence upstream of the AUG is expected to (i) strengthen initiation complex–mRNA interactions near the start codon, (ii) reduce leaky scanning by reinforcing sequence discrimination at the authentic AUG, and (iii) mitigate inhibitory features across diverse 5’-UTR contexts. We also anticipated full compatibility with m^1^Ψ-modified transcripts, as the Kozak consensus sequence lacks uridines and would be unaffected by this base substitution. To test this, we generated mRNAs containing 0–4 additional GCCACC motifs (1–5 total Kozak contexts) and assessed translation efficiency in vitro and in vivo to evaluate both generalizability and the predicted non-monotonic optimum in motif number and spacing.

## RESULTS

### Incorporation of additional Kozak consensus sequences in the 5’-UTR influences translation

In the context of mRNA therapeutics, native 5’ UTRs from host genes can provide regulatory compatibility and support robust expression; however, the AUG-proximal Kozak context in many endogenous transcripts is suboptimal, contributing to variability in translation efficiency (29). We therefore hypothesized that adding strong canonical Kozak motifs could enhance coding sequence translation without disrupting other regulatory functions of the 5’-UTR.

To test this hypothesis, we cloned varying numbers of the optimal Kozak motif (5’-GCCACC-3’) directly upstream of the ATG start codon in the luciferase coding sequence, while preserving the native Kozak motif in the 5’-UTR of the *APOA2* gene (APO). This approach was designed to maintain the regulatory functions of the UTRs as much as possible. Specifically, we introduced the Kozak motif in tandem 0, 1, 2, 3 or 4 times, resulting in mRNAs containing 1, 2, 3, 4, or 5 Kozak sequences in total, hereafter denoted as 1*k, 2*k, 3*k, 4*k, and 5*k, respectively (Figure 1A). Other mRNA elements, such as the poly(A) tail, coding sequence, and 3’-UTR, were kept constant. Specifically, the 3’-UTR was derived from the human β-globin gene (HBB) and duplicated, a configuration that has been reported to enhance translation (30, 31). In vitro-transcribed mRNAs were then transfected into HeLa cells, and luciferase activity was measured to evaluate translation efficiency.

**Figure 1.**
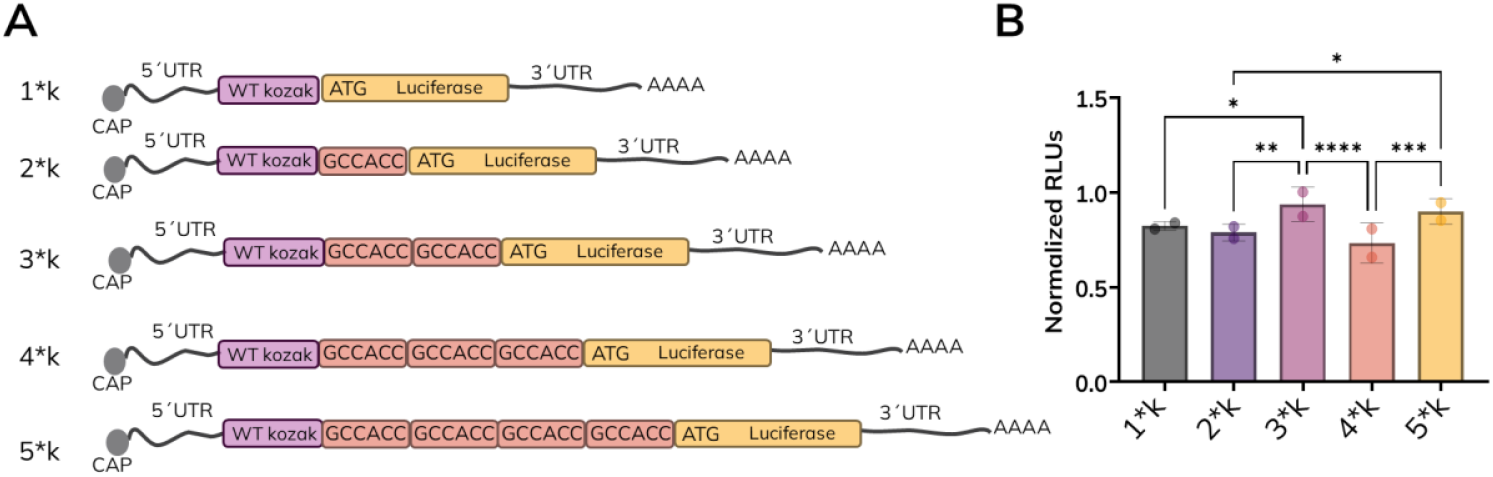
Overview of Kozak motif additions to mRNA constructs and their in vitro translation efficiency. **A)** Schematic representation of the mRNA constructs. The added canonical Kozak motifs (GCCACC) are highlighted in purple boxes, while the wild-type (WT) Kozak motifs are shown in pink and Luciferase coding sequence in yellow. The 5’-CAP structure is depicted as a grey circle, with the 5’and 3’untranslated regions (UTRs) represented by grey lines, and the polyadenine tail indicated by “AAAA”. **B)** Luciferase expression in HeLa cells was quantified as Relative Luminescence Units (RLU) for mRNAs containing the APO 5’-UTR with the indicated Kozak motif additions and the HBB 3’-UTRs and normalized to a control mRNA. Bars represent the mean values from two independent experiments; each performed in triplicate. Data points reflect the average of triplicates for each independent experiment. Statistical significance was assessed using a two-way ANOVA followed by Fisher’s LSD multiple comparison test. Asterisks indicate statistical significance between groups with the following p-values: * *p* ≤ 0.05, ** *p* ≤ 0.01, *** *p* ≤ 0.001, and **** *p* ≤ 0.0001.

The results showed that the addition of Kozak motifs did not directly correlate with increased translation efficiency (Figure 1B). Interestingly, the presence of three Kozak motifs (3*k) resulted in a statistically significant increase in translation compared to the mRNA containing the wild type Kozak motif (1*k) and to the groups with two or four motifs (2*k and 4*k). Moreover, an increase in translation was observed when odd numbers of Kozak motifs were present (3*k and 5*k), although the effect of 5k was not statistically significant.

### Validation of Kozak sequence addition with diverse 5’ and 3’-UTRs

To further validate the impact of adding Kozak sequences on mRNA translation efficiency, we extended our experiments to include the widely used human β-globin (HBB) 5’-UTR while keeping the duplicated HBB 3’-UTR constant. Specifically, we compared 1*k with 2*k and 3*k (Figure 2A) as 3*k modification demonstrated the highest translation efficiency (Figure 1B).

**Figure 2.**
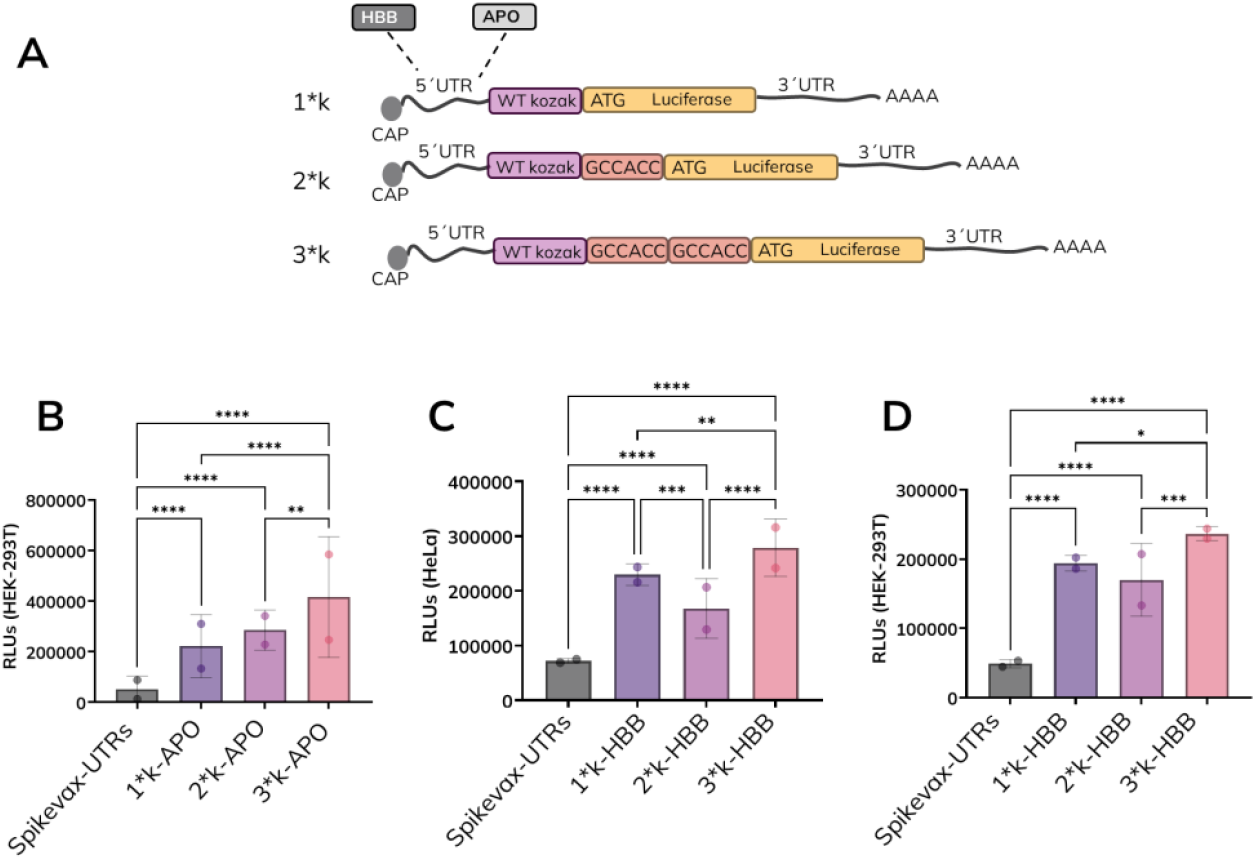
Effect of Kozak motif addition and 5’UTR variations on translation efficiency. **A)** Schematic representation of the mRNA constructs analyzed. The added canonical Kozak motifs (GCCACC) are in purple boxes, while the WT Kozak motifs are shown in pink and the Luciferase coding sequence in yellow. The 5’-CAP structure is represented by a grey circle, and the polyadenine tail is indicated as “AAAA.” The analyzed 5’-UTRs are labeled as HBB (human β-globin) in dark grey and APO (apolipoprotein) in light grey. **B)** Luciferase expression in HEK293T cells was quantified as Relative Luminescence Units (RLU) for mRNAs containing the 5’-UTRs of APO with the indicated Kozak motif additions. Luminescence in **C)** HeLa cells and **D)** HEK293T cells for mRNAs containing the 5’-UTRs of HBB with the indicated Kozak motif additions. Bars represent the mean values from two independent experiments; each performed in triplicate. Data points represent the average of triplicates for each independent experiment. Statistical significance was assessed using two-way ANOVA followed by Fisher’s LSD multiple comparison test. Asterisks indicate statistical significance between groups with the following p-values: * *p* ≤ 0.05, ** *p* ≤ 0.01, *** *p* ≤ 0.001, and **** *p* ≤ 0.0001.

We first confirmed if the performance of APO 5’-UTRs in HeLa cells (Figure 1B) was reproducible in HEK293T cells, a commonly used cell line for transfection studies (Figure 2B). Consistent with the results in HeLa cells, the incorporation of two additional Kozak sequences (3*k) resulted in a significant increase in luciferase expression compared to both the WT Kozak motif (1*k) and the single Kozak addition (2*k), demonstrating a similar enhancement in translation efficiency across both cell lines.

Next, we measured luciferase expression in HeLa (Figure 2C) and HEK293T (Figure 2D) cells using mRNAs containing HBB 5’-UTRs with 1*k, 2*k, or 3*k. In both cell lines, the 3*k modification resulted in a notable and statistically significant increase in luciferase expression compared to the WT Kozak motif, further supporting the utility of this approach.

Interestingly, we compared our UTR designs to those utilized in the commercial Spikevax mRNA vaccine against SARS-CoV-2. In all cases, regardless of whether additional Kozak motifs were included, our designs consistently outperformed the Spikevax UTRs, demonstrating superior translation efficiency.

Given that the 3’-UTR, in combination with the 5’-UTR, can significantly affect translation efficiency (32), we next investigated whether modifying the 3’-UTR to match the 5’-UTR, while avoiding sequence duplications, would influence the translation efficiency in constructs with added Kozak motifs (Figure 3A).

**Figure 3.**
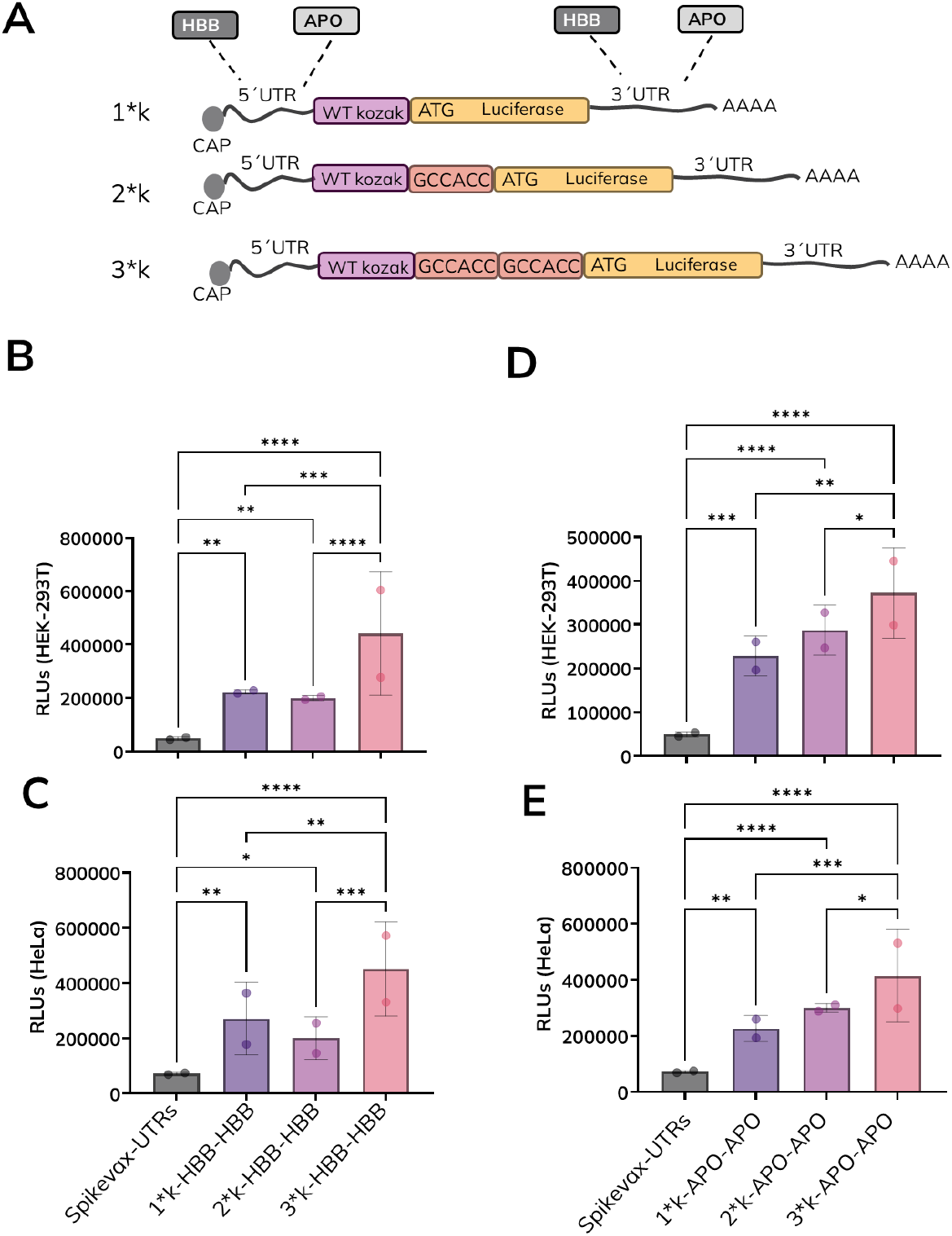
Effect of Kozak motif addition and 3’UTR variations on translation efficiency. **A)** Illustration of the mRNA constructs. Added GCCACC Kozak motifs are shown in purple, WT Kozak in pink, luciferase CDS in yellow, 5′ cap as a grey circle, poly(A) tail as “AAAA,” and UTRs as HBB (dark grey) or APO (light grey). **B)** Luciferase expression in HEK293T cells and **C)** HeLa cells was quantified as Relative Luminescence Units (RLU) for mRNAs containing the 5’-UTRs of HBB with the indicated Kozak motif additions and the 3’-UTRs of HBB. **D)** Luminescence in HEK293T and **E)** HeLa cells transfected with mRNAs containing the 5’-UTRs of APO with the indicated Kozak motif additions and the 3’-UTRs of APO. Bars represent the mean values from two independent experiments; each performed in triplicate. Data points represent the average of triplicates for each independent experiment. Statistical significance was assessed using two-way ANOVA followed by Fisher’s LSD multiple comparison test. Asterisks indicate statistical significance between groups with the following p-values: * p ≤ 0.05, ** p ≤ 0.01, *** p ≤ 0.001, and **** p ≤ 0.0001.

We first quantified luciferase expression in HEK293T (Figure 3B), and HeLa (Figure 3C) cells transfected with mRNA constructs containing the 5’-UTR and corresponding 3’-UTR of HBB. These mRNA constructs included the WT Kozak motif as well as variants with additional canonical Kozak sequences. In both cell lines, constructs with three Kozak sequences (3*k) consistently exhibited the highest levels of luciferase expression. A similar analysis was performed using mRNA constructs with the APO 5’-UTR and 3’-UTR in HEK293T (Figure 3D) and HeLa (Figure 3E) cells. Once again, the 3*k variant demonstrated the greatest enhancement in luciferase expression across both cell types, mirroring the results observed with the HBB constructs. Notably, in all tested conditions, our Kozak-optimized constructs consistently outperformed those incorporating the UTRs from the Spikevax mRNA vaccine. These findings clearly demonstrate that the addition of two Kozak motifs significantly enhances translation efficiency across various 5’ and 3’-UTR combinations in widely used HEK293T and HeLa cell lines.

### In vivo impact of Kozak motif additions on mRNA translation

To determine whether the gains observed in vitro translate in vivo, we formulated luciferase mRNAs bearing the APO 5′-UTR and a duplicated HBB 3′-UTR and introduced one or two additional Kozak cores immediately upstream of the AUG (yielding 2*k and 3*k designs) (Figure 4A). LNPs were prepared with the ionizable lipid SM-102 and administered intramuscularly to mice. Luciferase activity in the injected muscle was quantified 4 h post-dose (Figure 4B).

**Figure 4.**
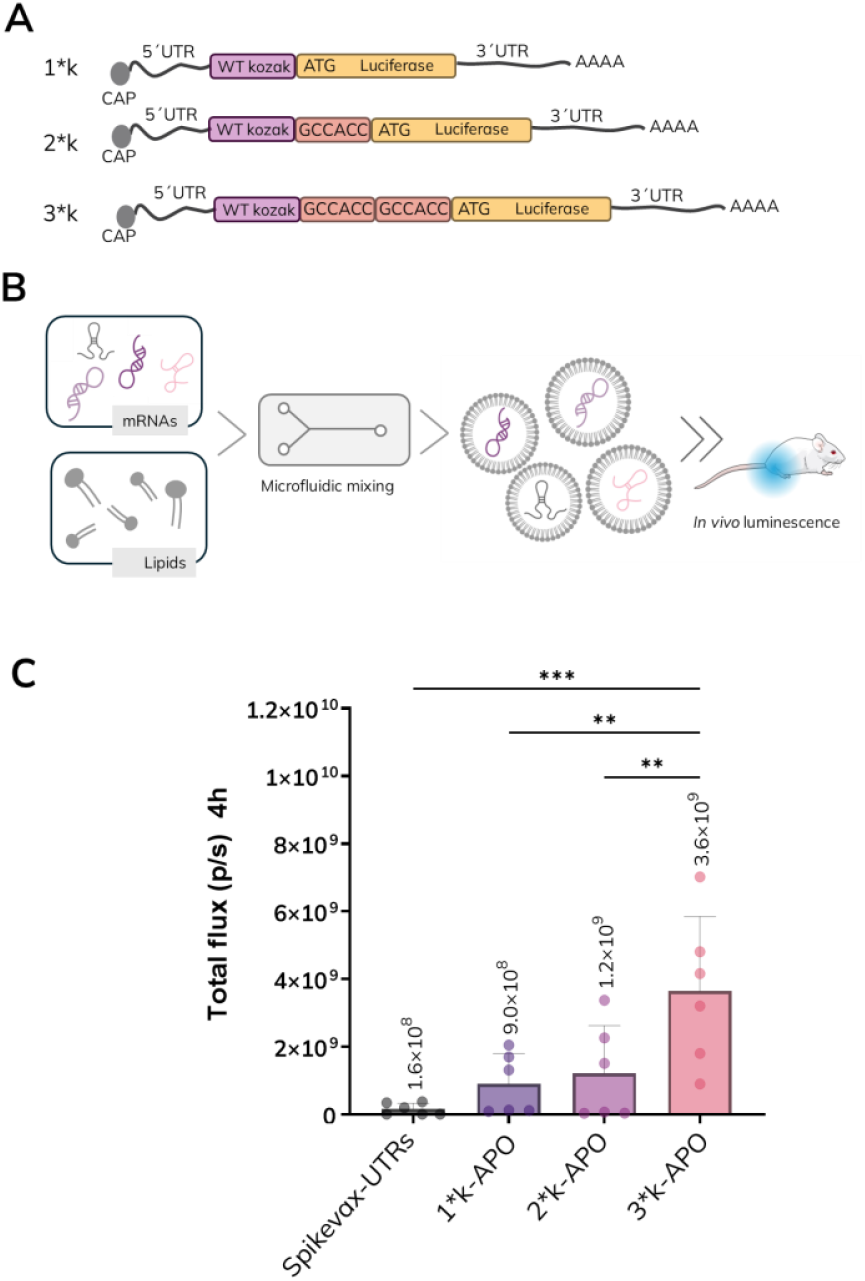
In vivo evaluation of tandem Kozak motif additions on mRNA translation efficiency. **A)** Diagram of the mRNA constructs analyzed. The added canonical Kozak motifs (GCCACC) are in purple boxes, while the WT Kozak motifs are shown in pink and the Luciferase coding sequence in yellow. The 5’-CAP structure is represented by a grey circle, and the polyadenine tail is indicated as “AAAA.” **B)** Workflow illustrating mRNA formulation into lipid nanoparticles (LNPs) followed by in vivo administration. **C)** Luciferase expression in mice for mRNAs containing the 5’-UTRs of APO with the indicated Kozak motif additions. Data are presented as mean ± SD from two independent experiments, each performed in triplicate. Data points represent each mice individual luminescence. Statistical significance was assessed using one-way ANOVA followed by Fisher’s LSD multiple comparison test. Asterisks indicate statistical significance between groups with the following p-values: ** p ≤ 0.01 and *** p ≤ 0.001.

Consistent with our cell-based assays, the 3*k variant was the clear top performer, producing a ∼4-fold increase in luminescence relative to the native context (1*k). Strikingly, when benchmarked against constructs carrying the 5′/3′-UTRs used in Spikevax, the 3*k-APO design delivered a ∼ 23-fold higher signal under matched dose and formulation. Moreover, 3*k construct exhibits higher inter-experimental consistency than the other designs (Figure 4C). This results further support the utility of Kozak motif addition for enhancing *in vivo* mRNA translation. These results not only validate the utility of Kozak motif addition for enhancing in vivo mRNA translation, but also demonstrate that a compact, easily implemented modification near the start codon can outperform full UTR designs offering a powerful and practical strategy for dose-sparing mRNA therapeutics.

## DISCUSSION

Our data show that strengthening the AUG-proximal context with short tandem repeats of the Kozak core (GCCACC) increases protein output in cells and in vivo, but with a non-proportional relationship to copy number. Across APO and HBB 5’-UTRs, three repeats (3*k) consistently outperformed the native sequence in HeLa and HEK293T cells, and enhanced luciferase expression ∼4-fold in mouse muscle following LNP delivery. Notably, the same design surpassed a licensed vaccine UTR benchmark by ∼23-fold, highlighting in vivo relevance.

These results support a simple, UTR-agnostic principle: tuning the local Kozak environment immediately upstream of the start codon can yield dose-sparing gains without re-engineering the entire UTR architecture. Because the GCCACC motif contains no uridines, it is inherently compatible with m^1^Ψ-modified transcripts used in therapeutic manufacturing pipelines.

Authorized COVID-19 mRNA vaccines illustrate that 5’-UTR design choices meaningfully differ across products. Public regulatory documentation for BNT162b2 specifies that its 5’-UTR derives from human α-globin with an optimized Kozak sequence (“hAg-Kozak”), explicitly chosen to increase translational efficiency (33). In contrast, sequence analyses of the Spikevax vaccine (mRNA-1273) reveal the use of a synthetic “V1-UTR”, composed of two elements. This design includes secondary a GC-rich region immediately upstream of an ACCAUG context thereby, introducing secondary structure flanking the start codon, an approach that may stabilize mRNA while potentially slowing ribosomal scanning in some settings (34). Notably, our optimized UTR designs yielded substantially higher translation than the UTRs used in Spikevax, an improvement we attribute primarily to strengthening the AUG-proximal context, while acknowledging that other variables (e.g., UTR length and secondary structure or 3’-UTR pairing) may also contribute.

By contrast, emerging comparative studies in infectious-disease models emphasize that the 5’-UTR can be a dominant driver of outcome and that “one-size-fits-all” solutions are unlikely. For mRNA tuberculosis vaccines, replacing the mRNA-1273 5’-UTR with the adenoviral tripartite leader increased luciferase expression in dendritic cells and improved vaccine efficacy in mice, while effects were cell-type specific (e.g., absent in THP-1) (35). Beyond single elements, combinatorial screens show that pairing of 5’--and 3’-UTRs strongly modulates translation and stability, reinforcing the need to consider the UTR set as a system rather than independent parts (32). Our observation that 3*k improves expression across multiple 5’/3’-UTR pairings is therefore encouraging, yet it does not obviate careful selection of distal UTR features for specific tissues and therapeutic indications.

Regarding why three Kozak cores act better than two or four we hypothesize that this may represent an optimal balance between sequence signal-to-noise contrast, kinetic sampling efficiency, and structural accessibility during 43S pre-initiation complex (PIC) scanning. A GC-rich ramp of approximately 18 nucleotides could enhance signal-to-noise at the cognate AUG, thereby reducing leaky scanning while avoiding premature initiation at near-cognate codons (24). This length may also function as a kinetic “landing zone”, allowing the PIC to decelerate and engage the AUG more effectively without incurring the dwell-time penalties or excessive helicase load with longer ramps. In addition, three repeats may maintain an unstructured window immediately upstream of the AUG, whereas additional motifs could increase the probability of forming inhibitory microstructures.

Our current work used luciferase reporters in two immortalized cell lines and a short (4 h) intramuscular mouse model, providing a robust framework to assess the Kozak core (3*k) design. Even so, factors such as reporter type, expression window, tissue context (35), and delivery platform may influence UTR performance, and exploring these variables will be a natural next step. Future studies evaluating 3*k in primary human cells, across additional UTR backbones, varied coding sequences, and different LNP chemistries will further strengthen the external applicability of our findings. Likewise, mechanistic investigations, such as ribosome profiling, single-molecule scanning, and RNA structure probing, will help clarify whether the observed gains are primarily driven by kinetic or structural effects.

In conclusion, this work demonstrates that precise tuning of the AUG-proximal context is a powerful and versatile strategy for enhancing translation efficiency. A three-copy Kozak ramp achieves an optimal balance between strong start-site commitment and minimal scanning burden, providing a compact, easily integrated feature fully compatible with existing therapeutic mRNA manufacturing and delivery platforms.

## MATERIALS AND METHODS

### Plasmid DNA template design for mRNA production

The coding sequence of luciferase (GenBank: WP_212371658.1) was codon-optimized for human expression and cloned in-frame immediately downstream of the designated 5’ -UTRs into a pUC57-based plasmid. The plasmid contained, in 5’ to 3’ orientation: a T7 RNA polymerase promoter, an AGG trinucleotide for co-transcriptional capping, the selected 5’ UTR, the luciferase coding sequence, the corresponding 3’-UTR from either *APOA2* or *HBB* genes, a poly(A) tail of 100 adenines with a single guanosine interrupting at position 30, and a BspQI restriction site. Complete 5’- and 3’-UTR sequences are provided in Supplementary Table 1.

The DNA template mimicking the Spikevax vaccine was generated by cloning the available mRNA vaccine sequence (36) into a pUC-based plasmid under control of a T7 promoter, replacing the spike coding sequence with that of luciferase and using NotI restriction site instead of BspQI.

Gene synthesis, cloning, and plasmid preparation were performed by Genscript. Purified plasmids were subsequently used as templates for in vitro transcription.

### mRNA synthesis and purification

Each plasmid containing the DNA template of interest was digested with BspQI (HONGENE, ON-124) or NotI (NEB, R3189B), which cleaved the plasmid immediately downstream of the segment to be transcribed. The resulting linearized DNA was then purified using the Wizard® SV Gel and PCR Clean-Up System (Promega, A7270) according to the manufacturer’s protocol.

The purified linear DNA was subsequently utilized for mRNA production through in vitro transcription, employing T7 RNA polymerase. Transcription reactions were conducted at 37°C for 3 hours, using the following components in RNase-free double-distilled water: linear DNA template (50 μg/mL), T7 RNA polymerase (5000 U/mL; HONGENE, ON-004), RNase inhibitor (1000 U/mL; HONGENE, ON-039), inorganic pyrophosphatase (2 U/mL; HONGENE, ON-025), ATP (2,8 mg/mL; HONGENE, R1331), GTP (2,8 mg/mL; HONGENE, R2331), CTP (2,6 mg/mL; HONGENE, R3331), N1-Methylpseudouridine (2,5 mg/mL; HONGENE, R5-027) and m7G(5’)ppp(5’)(2’-OMeA)pG (4,6 mg/mL;HONGENE, ON-134).

Following a 3-hour incubation, DNase I (Hongene, ON-109) was added to the synthesized mRNA transcripts, and the reaction was incubated for an additional 15 minutes at 37°C. The crude RNA was then purified via affinity chromatography using a POROS Oligo (dT) 25 column (ThermoFisher). The purification process utilized Buffer A (50 mM disodium phosphate, 0.5 M NaCl, 5 mM EDTA, pH 7.0) and Buffer B (50 mM sodium dihydrogen phosphate, 5 mM EDTA, pH 7.0). The mRNA samples were initially diluted two-fold with Buffer A. The column was equilibrated with 100% Buffer A, loaded with the mRNA, washed with Buffer B, and eluted with double-deionized water. To ensure complete removal of Buffer B, the mRNA was washed using a 30 kDa Amicon filter and equilibrated by a ten-fold dilution in 10x citrate buffer at pH 6.5.

The mRNA concentration was determined by measuring the optical density at 260 nm, adjusted to a final concentration of 1 mg/mL, aliquoted, and stored at -80°C. For quality control, all mRNA samples were analyzed using automated electrophoresis (2100 Bioanalyzer, Agilent, G2938B) before being aliquoted and stored at -80°C until further use.

### In vitro mRNA transfection

The HeLa (DSMZ GmbH, ACC57) cell line was maintained in DMEM with high glucose (Merck, D6429), supplemented with 10% Fetal Bovine Serum (Sigma, F7524), 1% Penicillin-Streptomycin Solution (GibcoTM, 15140122), and 2 mM Glutamax (ThermoFisher, 35050038). The HepG2 (ATCC, HB-8065) cell line was cultured in RPMI 1640 (GibcoTM, 31870074) with the same supplements: 10% Fetal Bovine Serum (Sigma, F7524), 1% Penicillin-Streptomycin Solution (GibcoTM, 15140122), and 2 mM Glutamax (ThermoFisher, 35050038). Both cell lines were grown in 175 cm^2^ flasks.

The day prior to transfection, cells were detached from the flasks using trypsinization (ThermoFisher, 11590626) and then seeded into 96-well plates at a density of 1 × 104 cells per well.

For transfections using a commercial cationic lipid, the culture media was replaced with 90 μL of fresh media. A mixture of mRNA (100 ng/well) and Lipofectamine MessengerMAX™ (Invitrogen, 15397974; 0.2 μL/well) was pre-incubated in OptiMEM media. This mRNA-lipofectamine mixture was then added to the corresponding wells in triplicate, resulting in a final mRNA concentration of 100 ng/well. In alternative conditions, the mRNA-Lipofectamine mixture was diluted to half or a quarter of its concentration before being added to the cell culture, achieving final mRNA concentrations of 50 ng/well and 25 ng/well, respectively.

The cells, along with the mRNA-Lipofectamine MessengerMAX mixture were incubated for 24 hours at 37 °C in a 5% CO2-atmosphere.

### Luciferase activity quantification in vitro

Cells were lysed 24 hours post-transfection by incubating 100 μL of PBS-Triton 0.1% for 10 minutes. Subsequently, 98 μL of the cell lysate was transferred to an opaque 96-well white plate. To each well, 100 μL of buffered d-Luciferin (GoldBio, LUCK-100) in 100 mM Tris-HCl pH 7.8, 5 mM MgCl2, 250 μM CoA, and 150 μM ATP buffer was added, resulting in a final concentration of 150 μg/mL. Cells that had not been treated with any mRNA served as the negative control. Luminescence was measured after 5 minutes of incubation at room temperature using a FLUOstar Omega plate reader (BMG LABTECH).

### LNP preparation and characterization

LNP synthesis was carried out using a microfluidic approach. An ethanolic lipid mixture was prepared containing the commercial SM-102 ionizable lipid (BocSCI, Shirley, NY, USA), along with DOPE (Corden Pharma, Plankstadt, Germany), cholesterol (Merck, Rahway, NJ, USA), and DMG-PEG2000 (Cayman, Ann Arbor, MI, USA) in a molar ratio of 50:10:38.5:1.5. This lipid solution was mixed with an aqueous phase consisting of mRNA in 10 mM citrate buffer (pH 4), yielding an ionizable lipid-to-RNA weight ratio of 10:1. LNP assembly was performed on an INano™ microfluidic system operating at a total flow rate of 12 mL/min, with an aqueous-to-ethanol flow rate ratio of 3:1. Following formation, residual ethanol was removed by dialysis using a Pur-A-Lyzer™ Midi Dialysis Kit (Sigma-Aldrich, St. Louis, MO, USA).

For physicochemical characterization, average particle size, polydispersity index (PDI), and zeta potential were measured with a Malvern Zetasizer Advance Lab Blue Label (Malvern Instruments Ltd., Malvern, UK). Encapsulated mRNA concentration was determined using the Quant-IT® RiboGreen assay (Invitrogen, Waltham, MA, USA) according to the manufacturer’s instructions, and encapsulation efficiency was confirmed by agarose gel electrophoresis.

The final LNP preparations were adjusted to a concentration of 100 μg/mL mRNA and stored at 4°C until use in in vivo studies.

### Administration of LNP-mRNAs in mice

Female BALB/c mice (Janvier), 8–10 weeks old and weighing between 18 and 23 g, were allowed to acclimate to the experimental facility for 3–7 days after arrival. Animals were kept under standardized housing conditions with controlled temperature (20–24°C), relative humidity (50–70%), and light intensity (60 lux) under a 12 h light/dark cycle.

For in vivo assessment of Firefly Luciferase expression, lipid nanoparticles (LNPs) containing 1 μg of the designated RNA in a total volume of 30 μL were administered by intramuscular injection. Four hours post-injection, mice were anesthetized using 4% isoflurane for induction (via vaporizer) and maintained at 1.5% during the procedure. Subsequently, D-luciferin (Quimigen, Madrid, Spain) was delivered intraperitoneally at 150 mg/kg. Luminescence imaging was performed 10 minutes after substrate administration using the IVIS Lumina XRMS Imaging System with Living Image software (version 4.8.2).

## ACKNOWLEDGEMENTS

We sincerely thank all members of Certest Biotec, and in particular the Certest Pharma team, for their invaluable insights, expertise, and support, which greatly contributed to the success of this work. We also acknowledge the use of the “Servicios Científico-Técnicos” at CIBA (IACS–University of Zaragoza), with special appreciation for the collaboration of the Animal and Medical Imaging and Phenotyping facilities. The authors would like to thank the Government of Aragón (Spain) for its financial support through project IDMF/2021/0009, (Nuevas tecnologías para el diseño y obtención de vacunas de ARN en Aragón).

## AUTHOR CONTRIBUTIONS

E.B.: Conceptualization, Data curation, Formal analysis, Supervision, Investigation, Visualization, Writing—original draft. I.B.M: Conceptualization, Data curation, Writing— review and editing. V.L.: Investigation, Methodology, Writing—review and editing. C.M.: Investigation, Methodology, Writing—review and editing. A.T.C: Investigation, Methodology, Writing—review and editing. J.M.O: Conceptualization, Supervision, Funding acquisition, Writing—review and editing.

## Supplementary information

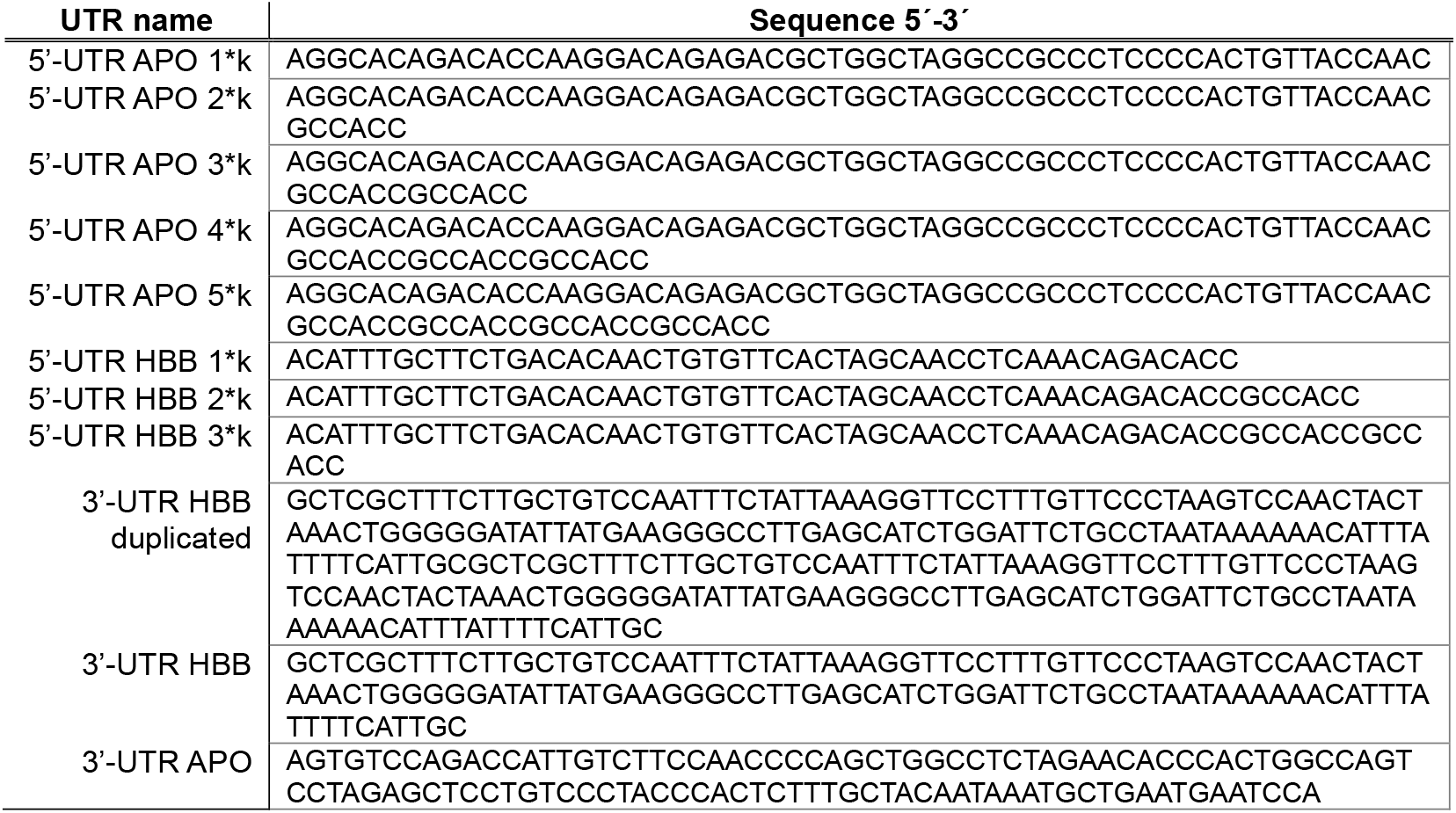

